# Reelin coordinates neuronal positioning and Müller glia scaffold maturation during retinal development

**DOI:** 10.64898/2026.08.16.745098

**Authors:** Priya Purohit, Simran Purohit, Yinuo Meng, William Cho, Francesca Telese, Dorota Skowronska-Krawczyk

## Abstract

Reelin is a secreted extracellular matrix protein that regulates neuronal migration and layer formation in the developing brain, yet its role in retinal development remains incompletely defined. Here, we investigated Reelin function in retinal lamination using wild-type and Reeler (*Reln^−/−^)* mice, combining stage-resolved RNA in situ hybridization, immunohistochemistry, and single-nucleus RNA sequencing. We show that *Reln* is dynamically expressed in ganglion cell layer and inner nuclear layer neurons during retinal development and persists in discrete adult neuronal populations. Loss of Reelin leads to widespread defects in retinal organization affecting both neurons and Müller glia. In *Reln*^−/−^ retinas, Müller glia exhibit reduced *Glul* positive extensions, indicating impaired glial scaffold maturation. Early-born neuronal populations are also disrupted, with altered spatial organization markers associated with retinal ganglion cell differentiation within the ganglion cell layer at postnatal day 9. Horizontal cells are significantly reduced with dorsal-predominant vulnerability, while cone photoreceptors are generated in normal numbers but show incomplete positioning within the outer nuclear layer. Together, these findings identify Reelin as a key regulator of retinal lamination that coordinates neuronal positioning with Müller glia morphogenesis, extending its canonical role in brain development to the vertebrate retina.

## INTRODUCTION

Reelin, encoded by the *Reln* gene, is a large secreted extracellular glycoprotein that plays a central role in regulating neuronal migration and positioning in laminated brain structures such as the cerebral cortex, hippocampus, and cerebellum. Reelin signaling is mediated through the lipoprotein receptors VLDLR and ApoER2, which activate the intracellular adaptor protein DAB1 and downstream pathways that regulate cytoskeletal dynamics, neuronal polarization, and the terminal positioning and detachment of migrating neurons from radial glial scaffolds^1,2,3,4^. Beyond its canonical role in lamination, Reelin also modulates dendritic maturation, spine development, and synaptic plasticity across multiple brain regions^4,5^. Importantly, Reelin signaling can also influence radial glial morphology, highlighting the potential role of Reelin to coordinate neuronal positioning with glial organization. Disruption of Reelin signaling has been associated with a range of neurodevelopmental and neuropsychiatric disorders, including lissencephaly, autism spectrum disorders, and schizophrenia, as well as broader cognitive impairments^6,7,8^.

In the visual system, components of the Reelin signaling pathway are expressed in the developing and mature retina, where they contribute to synaptic organization and neuronal connectivity. Reelin has been implicated in the establishment and maintenance of retinal synaptic architecture, particularly within plexiform layers, suggesting roles in circuit assembly in addition to its functions in the brain^9,10,11^. More recently, Reelin signaling was reported to protect retinal ganglion cells in an ischemia-reperfusion model, suggesting additional roles in retinal homeostasis^12^. However, how Reelin loss affects the laminar positioning of multiple retinal cell populations across development, and whether neuronal disorganization is accompanied by changes in Müller glial organization, remain incompletely defined.

Retinal development depends on the precise temporal and spatial coordination of cell proliferation, differentiation, migration, and final positioning. This process is orchestrated by retinal progenitor cells (RPCs) located in the neuroblastic layer (NBL), which progress through sequential competence states that bias them toward generating specific retinal cell types^13,14,15^. Early in development, RPCs predominantly generate retinal ganglion cells (RGCs), cone photoreceptors, horizontal cells, and subsets of amacrine cells, whereas later stages give rise to bipolar cells, rod photoreceptors, and Müller glia^16,17,18^. These developmental waves reflect dynamic transcriptional programs that integrate intrinsic genetic regulation with extrinsic signaling cues to determine cell fate and guide cells to appropriate laminar positions^13,14^. Proper apico–basal organization results in the formation of distinct retinal layers, including the outer nuclear layer (ONL: photoreceptors), inner nuclear layer (INL: bipolar, amacrine, horizontal cells, and Müller glia), and ganglion cell layer (GCL: RGCs), which are separated by the outer and inner plexiform layers (OPL and IPL) where synaptic connections are established^16,17^. Accurate neuronal migration and positioning are therefore essential prerequisites for functional circuit assembly and visual processing.

To determine the contribution of Reelin to retinal cell positioning and Müller glial organization, we combined stage-resolved RNAscope^®^ and immunohistochemistry with single-nucleus RNA sequencing (snRNA-seq). Using retinas from wild-type (WT, *Reln*^+/+^) and Reeler knockout mice (KO, *Reln*^−/−^), we mapped *Reln* expression and quantified the consequences of Reelin loss across developmental stages. Distinct retinal cell populations were examined using well-established lineage-specific markers. *Atoh7* (a basic helix–loop–helix transcription factor) was used to label RGC-competent progenitors^19,20^; differentiated RGCs were identified using *Brn3a* (POU-domain transcription factor), *Rbpms* (RNA-binding protein), and *Nefl* (neurofilament light chain)^21,22,23,24^. *Pax6* (paired-domain transcription factor) was used to label amacrine cell lineages^25,26^; *Arr3* (cone arrestin) marked maturing cone photoreceptors and their laminar localization within the ONL^27,28^; *Onecut1* (cut-like homeobox transcription factor) identified horizontal cells ^29,30^; and *Glul* transcript coding for glutamine synthetase was used as a marker for Müller glia^31,32^. This integrated approach enabled a lineage- and stage-resolved analysis of Reelin-associated retinal organization and allowed us to assess how Reelin deficiency affects the laminar distribution of neuronal population together with Müller-associated organization.

## RESULTS

### *Reln* Expression is Enriched in Ganglion Cell Layer and Inner Nuclear Layer Neurons

To establish the spatiotemporal pattern of *Reln* expression in the developing and mature retina, we performed fluorescent in situ hybridization (RNAscope^®^) at E12.5, E14.5, E17.5 and as 3-month-old retinas of *Reln^+/+^* mice.

At E12.5, Hoechst staining revealed a densely packed neuroblastic layer (NBL) without morphologically distinct retinal layers, consistent with ongoing proliferation (**Fig. 1A, B**). At this stage, *Reln* and *Glul* transcripts were broadly distributed throughout the NBL, consistent with the presence of both neuronal and glia progenitor-like populations in the undifferentiated retina. By E14.5, when the ganglion cell layer (GCL) first became morphologically apparent, *Reln* transcript was largely enriched in the emerging GCL, while *Glul* continued to be diffusely expressed throughout the retina. By E17.5, as the retinal structure expanded, *Reln* expression was concentrated in the GCL with additional puncta scattered in the forming inner nuclear layer (INL), while *Glul* expression persisted across the retina and was strongest near the ciliary margin zone, the most undifferentiated region of the retina.

**Figure 1.**
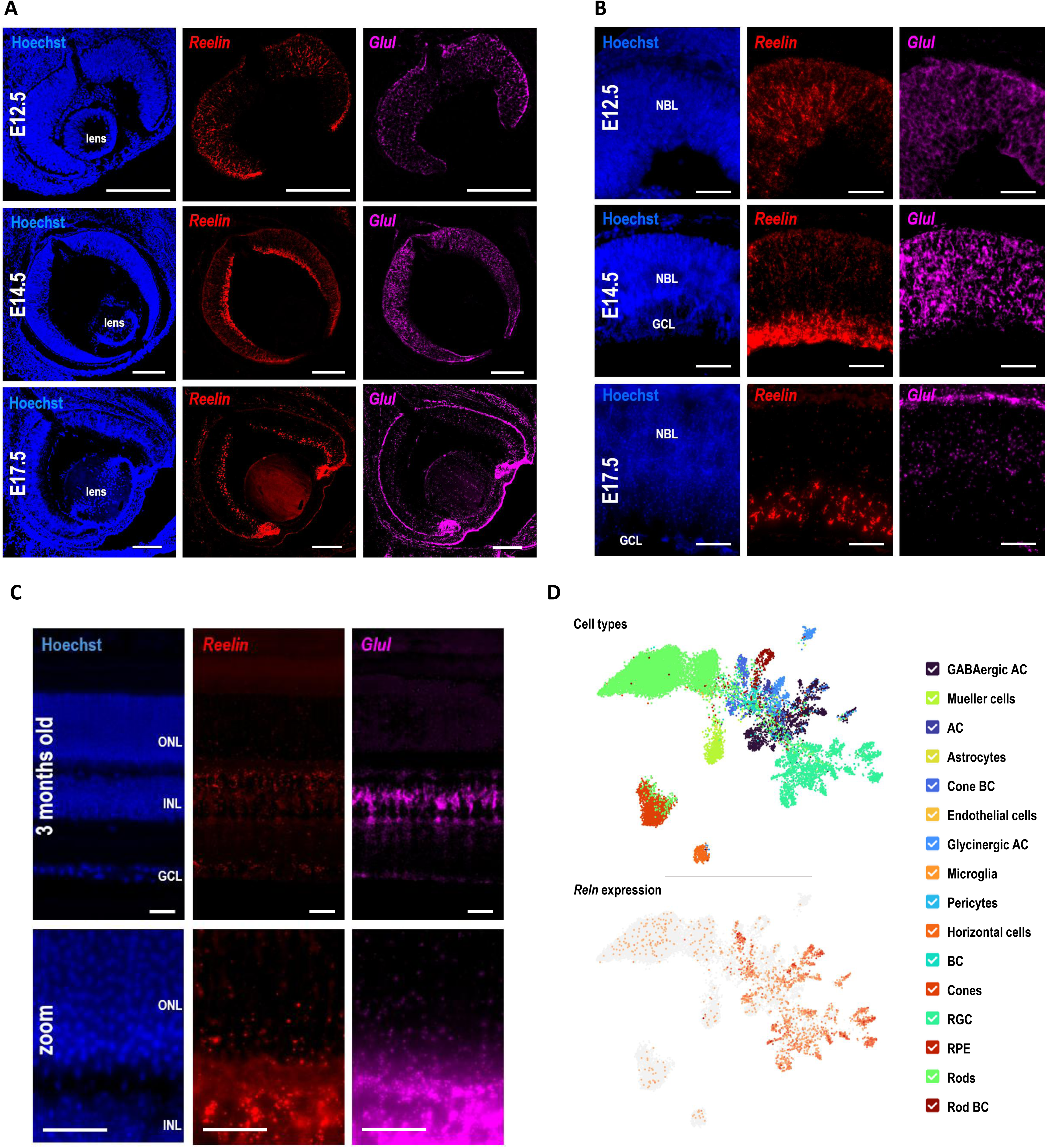
Reelin Expression During Mouse Retina Development. **(A)** Whole-eye sections at E12.5, E14.5, and E17.5 stained with Hoechst and RNAscope^®^ probes for *Reln* and *Glul*. Scale bar = 200 µm. NBL= neuroblastic layer, GCL = ganglion cell layer, INL = inner nuclear layer. **(B)** Higher-magnification views of (A) highlighting *Reln* enrichment in the GCL and the more diffuse *Glul* signal. Scale bar = 50 µm. **(C)** 3-month-old adult retina showing *Reln* signal localized to inner retinal neurons (GCL, INL) and *Glul* labeling Müller cell bodies and processes. Scale bar = 20 µm. ONL = outer nuclear layer. **(D)** UMAP from single-nucleus RNA-seq of 3-month-old retinas, with clusters labeled by annotated cell type (top) and *Reln* expression (bottom).

In the 3-month-old *Reln^+/+^* retina (**Fig. 1C**), *Reln* expression was confined to the INL and GCL. In contrast, *Glul* expression labeled Müller glia cell bodies in the INL, with radial processes spanning the outer nuclear layer (ONL), and their basal end feet forming a thin layer just above the GCL, consistent with canonical Müller glia morphology and marker expression^33,34^. Notably, the two signals showed minimal overlap, indicating that *Reln* is predominantly neuronal. Analysis of snRNA-seq data from 3-month-old retinas supported the predominantly neuronal distribution of Reln, with expression detected in RGCs, GABAergic amacrine cells (GABAergic AC), cone bipolar cells (Cone BC), horizontal cells, rod bipolar cells (Rod BC), and pericytes, and minimal expression in Müller glia cells marked by *Aqp4* (**Fig. 1D**, **Fig. S1**).

### Reelin Knockout Alters *Glul*-positive Müller Glia Organization

In the developing cortex, Reelin controls the detachment and final positioning of migrating neurons, and can also influence radial glia morphology, whereas Reeler (*Reln^−/−^)* mice show reduced radial-glial complexity^35^. We therefore asked whether loss of Reelin is associated with altered Müller glia organization in the adult retina. To address this question, we compared adult *Reln^+/+^* and *Reln^−/−^* retinas using RNAscope^®^ probes for *Reln* and *Glul* (**Fig. 2A–B**).

**Figure 2.**
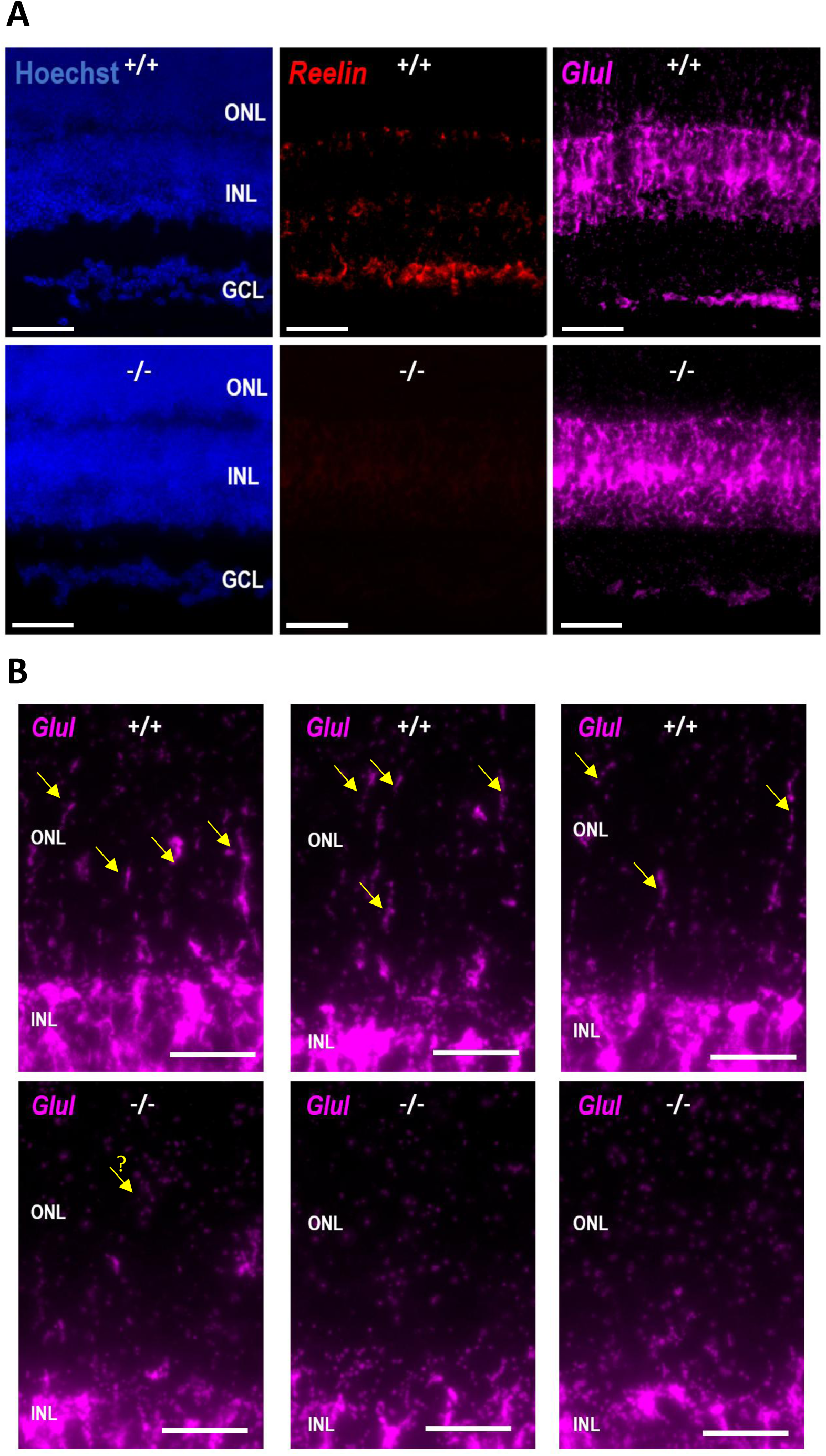
Reelin Loss Alters Müller Glia Architecture. **(A)** RNAscope^®^ labeling of *Reln* and *Glul* in adult wild-type (*Reln*^+/+^) and knockout (*Reln*^−/−^) retinas. Scale bars = 50 µm. GCL = ganglion cell layer, INL = inner nuclear layer, ONL = outer nuclear layer. The arrows are the processes of Müller glia. **(B)** Higher-magnification views of *Glul* labeling from (A) focusing on the ONL and INL. Scale bars = 250 µm. The arrows are the processes of Müller glia.

In the *Reln^+/+^* retinas, *Reln* transcripts were localized predominantly to the GCL and lower INL, with sparse puncta near the inner border of the ONL. In contrast, *Glul* signal was concentrated in the central INL with their basal endfeet forming a continuous layer just above the GCL, while their apical processes extending toward the ONL, consistent with the spatial distribution of Müller glia-assocatiated transcript labeling and a mature radial glial morphology.

As expected, *Reln* signal was absent in the *Reln^-/-^* knockout retinas. *Glul* signal remained detectable in the INL but showed fewer apparent radial extensions toward the ONL and a thinner, less continuous endfoot layer above the GCL (**Fig. 2A**, bottom). High-magnification views showed that *Reln*^+/+^ retinas displayed numerous *Glul*-positive apical processes traversing the ONL, whereas these processes were sparse or absent in *Reln*^−/−^ (**Fig. 2B**).

Together, these results demonstrate that Reelin loss is associated with altered spatial distribution of *Glul*-positive Müller glial labeling, suggesting that Reelin supports the full maturation of Müller glia radial architecture.

### Loss of Reelin Alters the Spatial Distribution of RGCs-Associated Markers

Because Reelin regulates the detachment and final positioning of migrating neurons in other laminated neural tissues, we next asked whether loss of Reelin alters the timing and spatial distribution of RGCs-associated markers. We examined *Atoh7* and *Nefl* as markers of RGC progenitors at E14.5.

In *Reln*^+/+^ retinas, the *Nefl* signal was broadly distributed across the developing retina, whereas in *Reln*^−/−^ retinas, it appeared more intense and concentrated in the central retina (**Fig. 3A**). *Atoh7* was detected in deeper layer regions in both genotypes, but, despite similar signal intensity, its distribution in *Reln*^−/−^ retinas appeared to partially occupy regions where *Nefl* signal was reduced, suggesting altered spatiotemporal coordination during RGC differentiation or delayed RGC maturation^36^.

**Figure 3.**
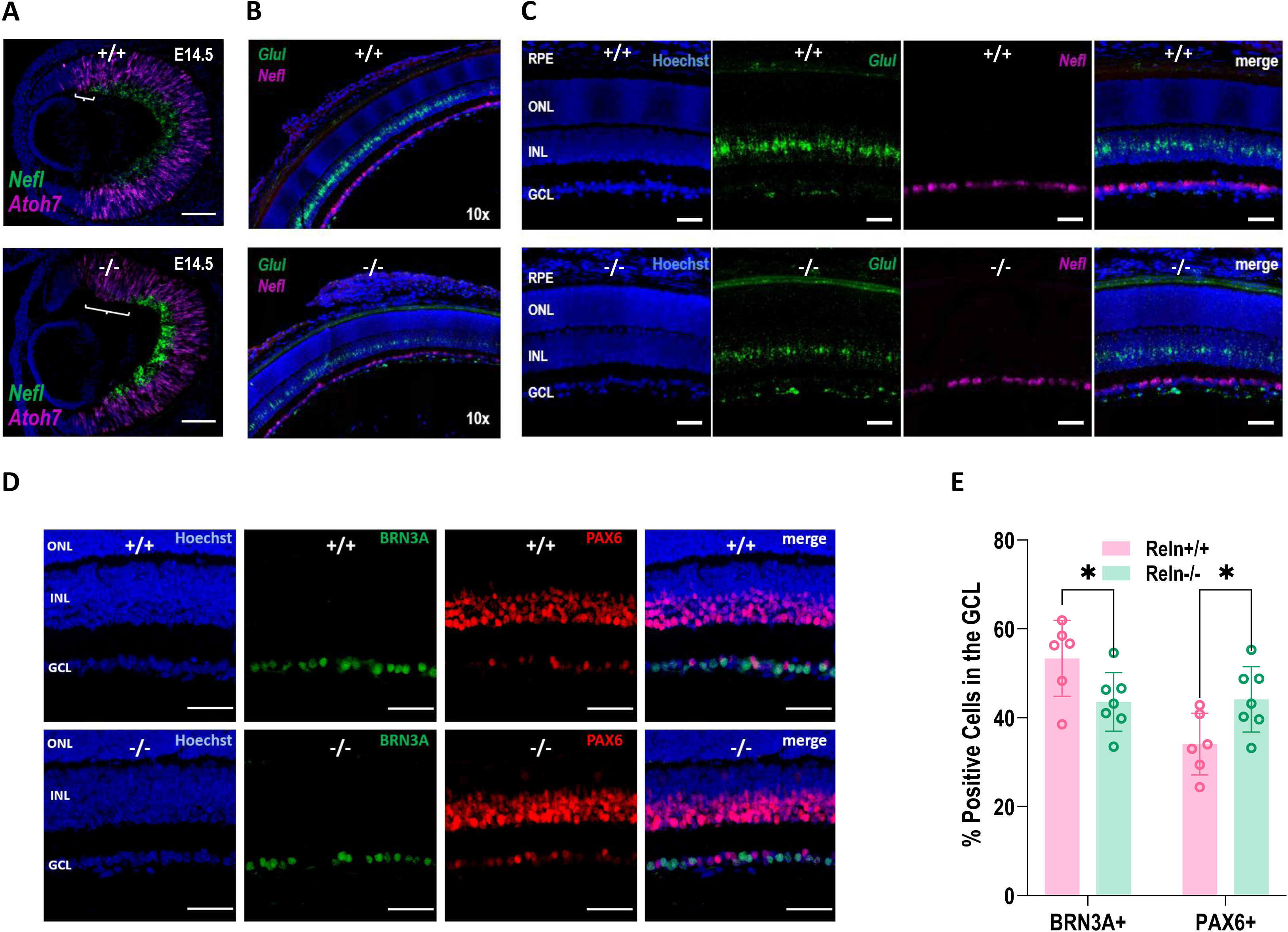
Loss of Reelin Alters RGC Distribution and Amacrine Cell Abundance. **(A)** RNAscope^®^ co-labeling of *Nefl* and *Atoh7* in E14.5 *Reln*^+/+^ and *Reln*^−/−^ retinas. Hoechst nuclear counterstain in blue. Scale bars = 200 µm. Brackets indicate the size difference in the region enriched in uncommitted precursors. **(B)** Merged 10x image of *Glul* and *Nefl* markers in a WT (*Reln*^+/+^) and MUT (*Reln*^-/-^) sample. **(C)** Higher-magnification views of *Glul* and *Nefl* at E14.5 *Reln*^+/+^ and *Reln*^−/−^ retinas. Scale bars = 50 µm. **(D)** Immunostaining for BRN3A and PAX6 in P9 retinas from *Reln*^+/+^ and *Reln*^−/−^ mice. Hoechst counterstain in blue. Scale bars = 100 µm. **(E)** Quantification of BRN3A+ and PAX6+ cells as a percentage of all GCL nuclei in *Reln*^+/+^ and *Reln*^−/−^ retinas (mean ± SEM; \**p* < 0.05, unpaired *t*-test).

We next assessed *Glul* and *Nefl* localization at E14.5 (Fig. 3B, C). Compared to *Reln*^+/+^ retinas, *Reln*^-/-^ retinas showed an altered distribution of *Glul* signal, with relatively more labeling near the developing GCL and less in the presumptive INL. *Nefl* signal showed similar results, with a slightly increased signal in the GCL in the *Reln^-/-^* sample compared to *Reln^+/+^*, suggesting premature or excessive accumulation of RGCs at this layer.

To determine whether altered cellular distribution persisted after birth, we immunostained for BRN3A and PAX6, markers of RGC and amacrine cells, respectively, in postnatal day 9 (P9) (**Fig. 3D-E**). The percentage of BRN3A+ cells were significantly reduced within the GCL of *Reln^-/-^* retinas compared to *Reln^+/+^* controls (**Fig. 3E**), whereas the percentage of PAX6+ cells were significantly increased. BRN3A+ cells were also observed in the INL of the *Reln^-/-^* samples, consistent with altered RGCs positioning and incomplete migration of amacrine cells.

Together, these data show that Reelin deficiency perturbs the normal spatial distribution of RGCs, with fewer BRN3A+ RGCs and a relative expansion of PAX6+ amacrine cells in the GCL.

### Reelin Deficiency Reduces *Onecut1*^+^ Horizontal Cells and Biases their Regional Distribution

Horizontal cells are early-born neurons that organize synaptic architecture at the OPL and might be particularly vulnerable to migration defects in the absence of Reelin. To determine whether Reelin loss affects horizontal cells, we used *Onecut1* as a marker and quantified their abundance and laminar position in *Reln*^+/+^ and *Reln*^−/−^ retinas (**Fig. 4A–B; Fig. S2**). *Onecut1* is essential for horizontal cell specification and survival^29^.

**Figure 4.**
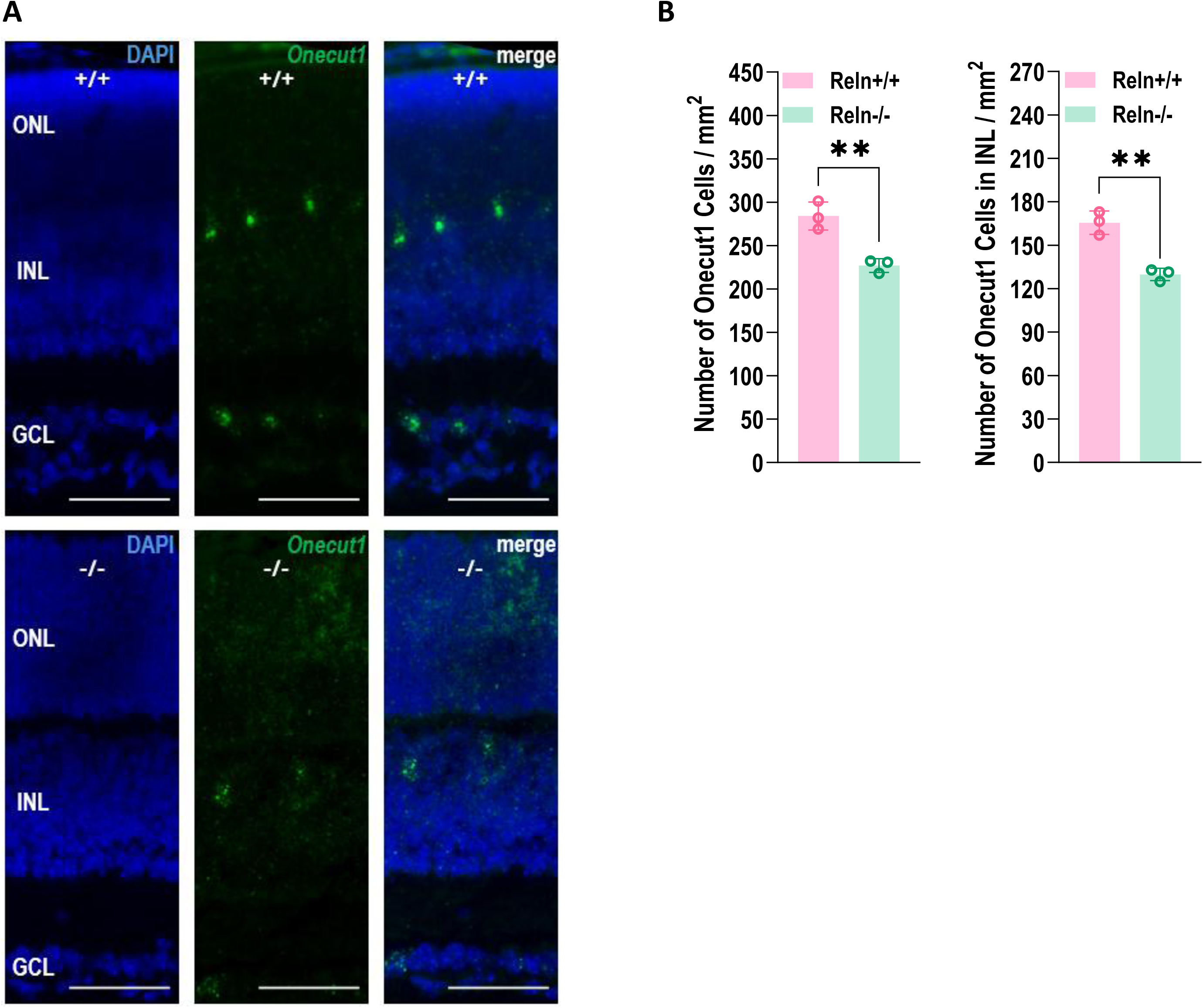
Reelin Deficiency Reduces *Onecut1*⁺ Horizontal Cells and Alters Outer-Retinal Organization. **(A)** Representative retinal sections from wild-type (*Reln*⁺^/^⁺) and mutant (*Reln*⁻^/^⁻) mice showing *Onecut1* (green) detected by RNAscope^®^ with DAPI (blue). Layers: GCL, INL, ONL. Scale bars = 50 µm. GCL = ganglion cell layer, INL = inner nuclear layer, ONL = outer nuclear layer. **(B)** Quantification of *Onecut1*^+^ cells: total cells per mm² (left) and INL-restricted cells per mm² (right) in *Reln*⁺^/^⁺ vs *Reln*^-/-^ retinas. Bars show mean ± SEM; individual dots denote individual fields, averaged per animal for statistics (n = 6 fields from 3 animals per genotype). Significance: *p < 0.01 (two-tailed t-test).

In *Reln*^+/+^ retinas, *Onecut1*^+^ horizontal cells formed a distinct band in the INL (**Fig. 4A**). In contrast, *Reln*^−/−^ retinas showed reduced *Onecut1* labeling (**Fig. 4A**), and quantification revealed a significant reduction in both the total number of *Onecut1*^+^ cells and the number within the INL compared to *Reln^+/+^* samples (**Fig. 4B**). We detected fewer *Onecut1^+^* cells also in the GCL (**Fig. S2A–B**). In regional analyses, the numerical reductions were more pronounced dorsally than ventrally (**Fig. S2B**).

Together, these results demonstrate reduced abundance and alter laminar allocation of *Onecut1^+^* cells within the INL in *Reln*^-/-^ retinas, with a stronger impact in the dorsal retina.

### Reelin Deficiency Alters Final Cone Positioning

Because cones, together with ganglion and horizontal cells, are among the cell types that develop early in the retina^37^, we next asked whether Reelin deficiency alters their abundance and their final positioning. We labeled cones with a probe targeting *Arr3* and quantified both their number and spatial distribution in *Reln*^+/+^ and *Reln*^−/−^ retinas (**Fig. 5A-C**).

**Figure 5.**
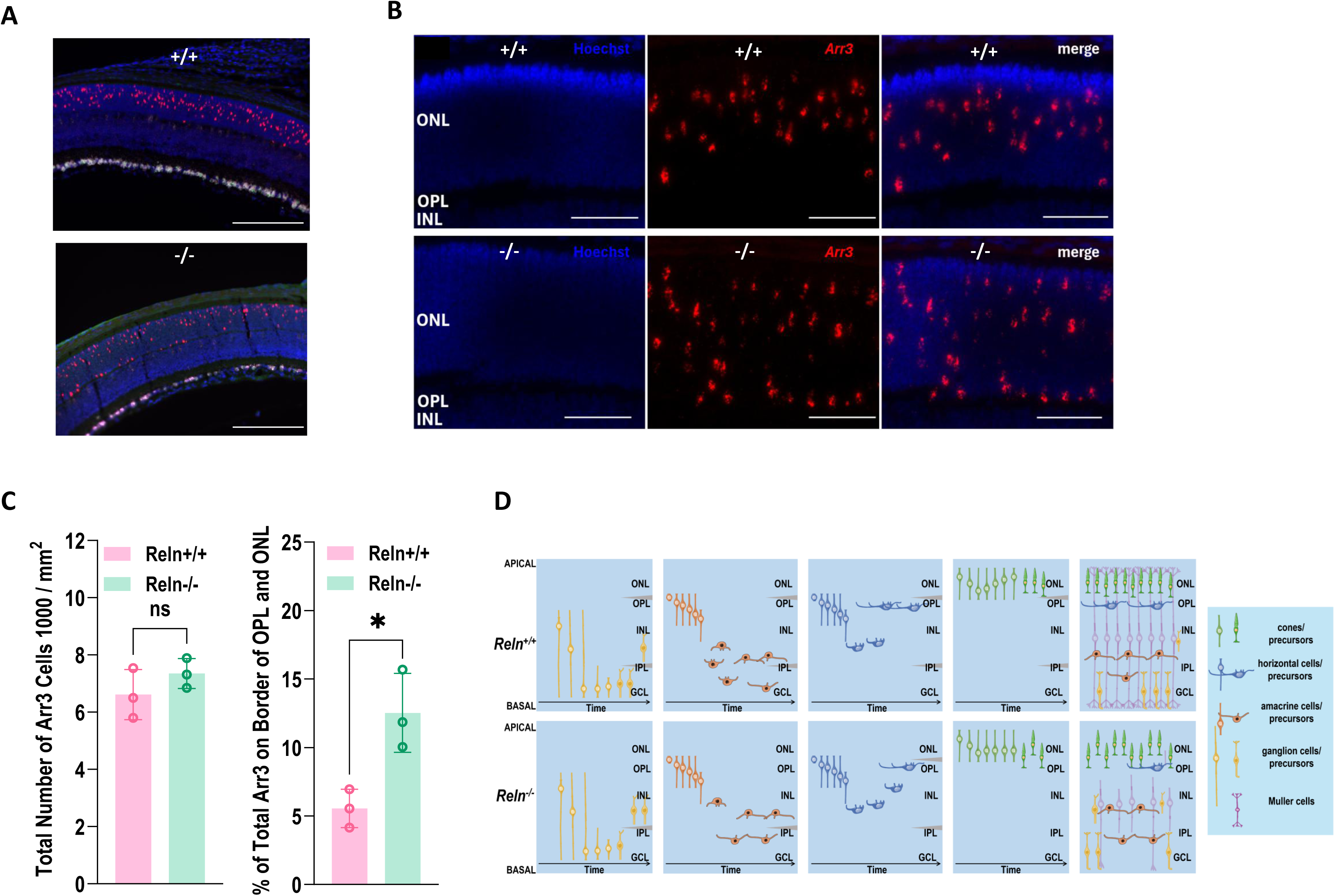
Reelin Deficiency Results in Stalled Cone Migration. **(A)** Low-magnification RNAscope^®^ images showing *Arr3*+ cones and *RBPMS*+ RGCs in *Reln*^+/+^ and *Reln*^−/−^ retinas. Hoechst nuclear counterstain in blue. Scale bars = 200 µm. **(B)** Higher-magnification images of *Arr3*+ cones with Hoechst nuclear counterstain in *Reln*^+/+^ and *Reln*^−/−^ retinas. Scale bars = 50 µm. **(C)** Quantification of *Arr3*+ cones per mm² (left) shows no difference between genotypes, but the percentage of cones located at the OPL/ONL border is significantly increased in *Reln*^−/−^ retinas (right). Bars represent mean ± SEM. Significance: \**p* < 0.05, unpaired *t*-test. **(D)** Working model summarizing Reelin-dependent lamination: in *Reln*⁻^/^⁻ retinas, cones do not fully migrate to the ONL and there are fewer horizontal cells, amacrine cells, retinal ganglion cells, and Müller glia cells in their correct respective locations.

Across genotypes, there was no significant change in the number of cones in the ONL (**Fig. 5C**). However, there were significantly higher percentages of *Arr3* positive cells positioned at the border between the outer plexiform layer (OPL) and the ONL in the *Reln^-/-^* samples compared with controls (**Fig. 5B-C**). This positional phenotype was observed across three independent biological replicate sets and in both dorsal and ventral retinal regions (**Supplementary Fig. 3**).

Together, these results demonstrate that Reelin deficiency does not substantially alter total cone abundance but is associated with incomplete final cone positioning at the OPL/ONL border.

## DISCUSSION

Our findings identify Reelin as an important regulator of retinal laminar organization. Using stage-resolved RNA-FISH, postnatal immunostaining, and adult single-nucleus RNA-seq, we found that *Reln* transcripts are enriched in neuronal population of the GCL, lower INL, and inner ONL, and that Reelin deficiency is associated with altered positioning or representation of several early-born retinal cell populations. Loss of Reelin resulted in truncated Müller glia processes, mispositioned RGCs, a dorsal-predominant reduction in *Onecut1*^+^ horizontal cells, and stalled cone migration. These findings extend the established role of Reelin in neuronal positioning in laminated brain structures to retinal development^1,2,3^, while also raising the possibility that Reelin influences neuronal-glial organization in the retina.

The effects of Reelin deficiency on Müller glia were particularly striking given the predominantly neuronal localization of *Reln* transcripts. Müller glia provide structural and metabolic support in the retina, guide late-born neuronal migration, and maintain synaptic homeostasis^38^. At E14.5, *Glul* expression in *Reln*^−/−^ retinas appeared redistributed toward the developing GCL (**Fig. 3B**), consistent with an early positional bias rather than a generalized reduction in Müller-lineage signal. In adulthood, *Reln*^−/−^ retinas exhibited fewer *Glul*-positive processes extending through the ONL together with a thinner and less continuous endfoot layer adjacent to the GCL (**Fig. 2**), suggesting incomplete elaboration of Müller glia morphology. Although Reelin has been best characterized for its neuron-autonomous role in promoting neuronal detachment from radial glial fibers, previous studies have also demonstrated that it can directly influence radial glial maturation^35^. Our observations therefore suggest that in the retina Reelin may contribute not only to neuronal positioning but also to the development and maintenance of the radial scaffold that supports retinal organization. This convergence with the role of Reelin in regulating radial glial complexity in the cerebral cortex raises the possibility that Reelin signaling promotes glial scaffold maturation as a common mechanism across laminated neuroepithelia^4,35^.

Consistent with this interpretation, loss of Reelin disrupted the normal spatial organization of early-born retinal neurons. During cortical development, Reelin deficiency delays neurogenesis and impairs neuronal migration^39^. Strikingly, in the retina, loss of Reelin resulted in higher levels of central *Nefl* expression, suggesting premature differentiation of RGCs. At the same time, *Atoh7*-positive progenitors occupied broader regions at the retinal margins where *Nefl* labeling was absent compared to the wild-type retina, suggesting altered coordination between RGC differentiation and positioning. These findings align with the established mechanism of RGC differentiation^40^, in which progenitors undergo a prolonged penultimate cell cycle followed by a final round of interkinetic nuclear migration, with nuclei returning to the apical surface for terminal mitosis, before newly born RGCs migrate basally and differentiate. Our data suggest that Reelin is required for the proper execution of this final nuclear migration and that its deficiency causes precocious differentiation of RGC precursors. Consequently, by postnatal day 9, these embryonic alterations were accompanied by fewer BRN3A+ retinal ganglion cells and a relative increase in PAX6+ amacrine cells within the GCL, echoing Reelin’s canonical role in cortical lamination^1^. These findings are consistent with the possibility that Reelin functions during a narrow developmental window, coincident with final interkinetic nuclear migration and early RGC differentiation.

Horizontal cells appeared especially sensitive to the absence of Reelin. *Reln*^−/−^ retinas exhibited fewer *Onecut1*^+^ horizontal cells overall and within the INL, with a more pronounced reduction in the dorsal retina. Because *Onecut1* is required for horizontal-cell specification and survival^29^, the observed phenotype may reflect impaired maintenance, altered allocation, or increased vulnerability of this population in the absence of Reelin signaling. The dorsal predominance of this effect further suggests that Reelin-dependent processes may interact with regional patterning mechanisms that establish retinal identity. Signaling pathways involved in dorsal–ventral patterning, including BMP, Wnt, and Vax2 networks, represent plausible candidates for such interactions and warrant future investigation^26,14^.

The effects of Reelin loss extended beyond the inner retina to additional early-born neuronal populations. Our data show that cone photoreceptors were generated in normal numbers in *Reln*^−/−^ retinas, yet a greater proportion remained positioned near the OPL/ONL border, indicating that Reelin is dispensable for cone genesis but facilitates efficient completion of cone positioning into the ONL. Mechanistically, previous studies have shown that Reelin’s stabilization of the actin cytoskeleton allows for the migration and nuclear positioning of cone nuclei to their proper location^41,42^. Specifically, signaling for this process is transmitted through apolipoprotein E receptor 2 (ApoER2) and very low density lipoprotein receptor (VLDLR) by orchestrating actin cytoskeleton and microtubule dynamics to move the nucleus within the cell. Interestingly, the pattern of cone localization in mutant retinas suggests defective Reelin/VLDLR signaling involving distal leading processes and the terminal positioning phase in layering^41,43^.

Several limitations should be acknowledged. While our data establish a requirement for Reelin in retinal lamination, the cellular mechanisms remain to be defined. Given the expression of canonical Reelin receptors (ApoER2/VLDLR) in retinal neurons, it is likely that Reelin acts through conserved DAB1-dependent signaling; however, whether this pathway operates cell-autonomously in each affected population or through secondary tissue-level effects remains unresolved. In addition, while our data consistently demonstrates altered positioning and distribution of multiple retinal cell types, they do not distinguish whether these phenotypes arise from defects in migration, maturation, survival, or combinations of these processes. Furthermore, our data suggest a previously underappreciated link between neuronal positioning and Müller glia morphogenesis; however, the observed Müller glia defects may arise either from direct Reelin signaling in Müller cells or indirectly through altered neuronal organization and feedback interactions. Future conditional and lineage-specific approaches will be required to distinguish these possibilities.

In summary, our findings identify Reelin as an important organizer of retinal development that coordinates neuronal positioning with Müller glia morphogenesis across multiple early-born neuronal populations. Building on prior studies implicating Reelin signaling in retinal synaptic organization and neuronal survival, our results extend its role to the coordinated establishment of retinal lamination during development. By integrating glial scaffold formation with neuronal allocation across retinal layers, Reelin contributes to the establishment of normal retinal architecture and broadens the developmental functions of this conserved signaling pathway beyond its well-established roles in the cerebral cortex.

## METHODS AND MATERIALS

### Animals

All experimental procedures were approved by the institutional animal care and use committee at the University of California, San Diego. Mice were housed in groups of 3-4 per cage and maintained on a 12-hour light/12-hour dark cycle with food and water ad libitum. *Reln^-/-^* mice were bred in-house from the B6C3Fe a/a-Relnrl/J line (The Jackson Laboratory, strain #000235). This strain is isogenic, having been backcrossed to the C57B6/J genetic background for over 20 generations. For postnatal timepoints, we used mice that were P7, P9 or 3 months old; for embryonic timepoints, we used 12.5, 14.5, and 17.5 day old embryonic stages.

### RNAscope^®^ Procedure

Following euthanasia and enucleation, the eyeballs were sliced to obtain a cryo-sectioned sample of the retina. In situ hybridization was performed using the RNAscope^®^ Multiplex Fluorescent Assay v2 (ACD Diagnostics). The probes used were designed by the manufacturer (see table). We followed the manual’s instructions with slight modifications as follows. Fresh frozen histologic sections of mouse eyes were pretreated using hydrogen peroxide and target retrieval reagents. The samples were incubated with Proteinase K for 40 seconds and boiled for 5 minutes. Probes were detected with TSA Plus® Fluorophores fluorescein: GFP, cyanine 3, and cyanine 5. The samples were then counterstained with either DAPI or Hoechst. Sections were mounted with Prolong Gold Antifade (Thermo Fisher) and imaged (Keyence BZ-X700)^44,45^.

| **RNAscope® Probe** | **Catalog Number** |
| --- | --- |
| Mm-*Atoh7*-01-C1 | 527231 |
| Mm-*Reelin*-C1 | 405981 |
| Mm-*RBPMS*-C2 | 527231-C2 |
| Mm-*Arr3*-C2 | 486551-C2 |
| Mm-*Nefl*-C3 | 493671-C3 |
| Mm-*Glul*-C3 | 426231-C3 |

### Single-nucleus RNA-seq

Sn-RNAseq analysis was performed as previously described (ref). In brief, retinas (n=3) were dissected and snap-frozen in liquid nitrogen. Deep frozen samples were processed according to the Illumina “Isolation of Nuclei for Single Cell RNA Sequencing & Tissues for Single Cell RNA Sequencing” procedure and subjected to the snRNAseq following the Illumina protocol.

#### Quality control, integration and clustering

91bp sequencing reads were generated by Illumina sequencing. Reads were mapped to the and counts at features were identified using the Cell Ranger for Single Cell Gene Expression software with default parameters (mkfastq and count functions). Quality control was performed, and cellular barcodes that matched the following three criteria were kept: number of unique molecular identifiers (UMIs) within three median absolute deviations (MADs) of the population median, number of expressed genes within three MADs of the population median, and the percentage of reads mapping to mitochondrial genes under 15%. Expression of approximately 1000 genes per nucleus was detected in each sample. Downstream analysis was performed using the Seurat (v4.3.0) R package.

### Immunohistochemistry

Following euthanasia, eyes were enucleated and fixed in 4% paraformaldehyde (PFA) in PBS (Affymetrix) for 1 hour and subsequently transferred to PBS. The eyes were then dissected, the retinas flat-mounted on microscope slides, and immunostained using a standard sandwich assay with anti-BRN3A antibodies (Millipore, MAB1585), anti-PAX6 antibodies (Millipore, AB2237) and secondary AlexaFluor 555 anti-mouse (Invitrogen, A32727). The samples were then counterstained with either DAPI or Hoechst. Mounted samples (Fluoromount, Southern Biotech 0100-01) were imaged in the fluorescent microscope at 20x magnification (Biorevo BZ-X700, Keyence)^45,46^.

### Microscope Quantification Imaging and Data

#### Microscope Imaging

Immunostaining and RNAscope^®^ procedures were completed to analyze the results of Reelin on the proteins and gene sets. For PAX6 and BRN3A, four total images were taken per sample using the x40 lens. For *Arr3* and *Onecut1*, three to four and four total images were taken per sample using the x20 lens, respectively. Folded or cut regions were excluded.

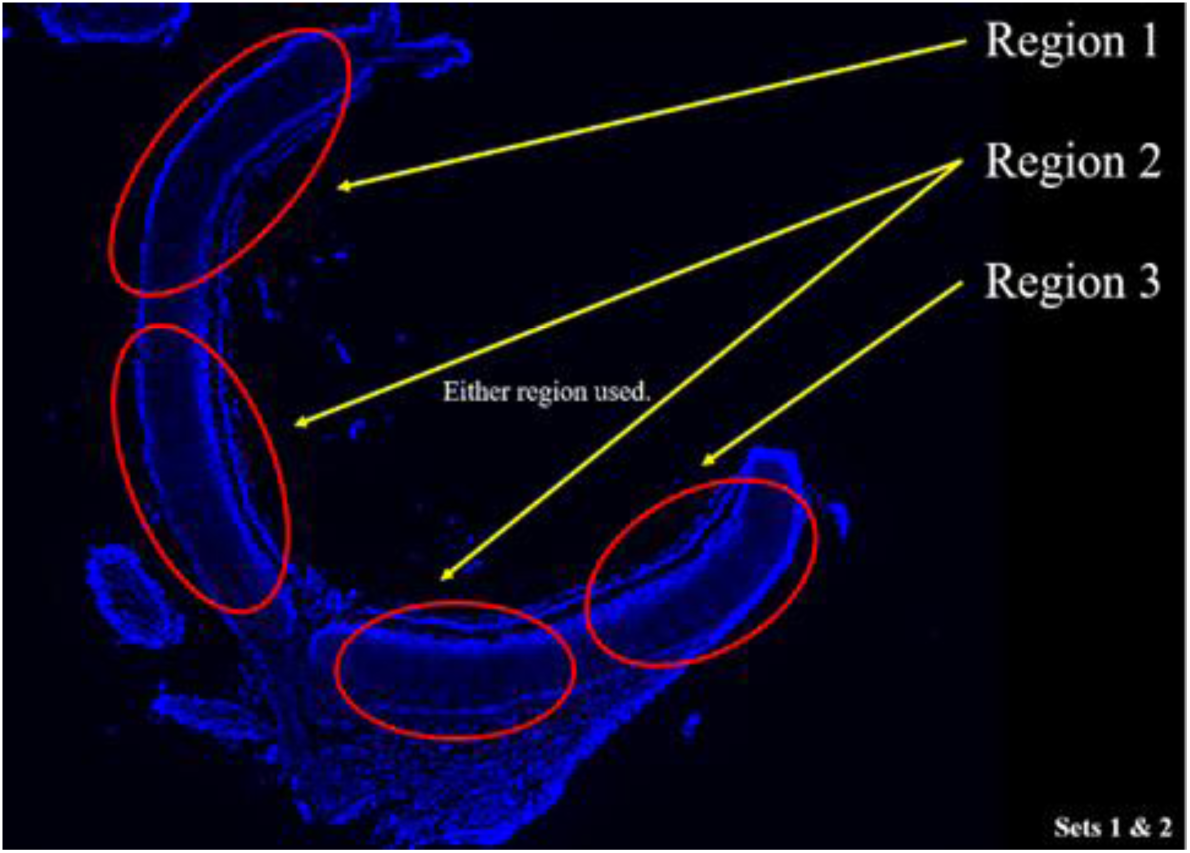

*Arr3* Microscope Imaging: The above image is a visual representation of each region in Sets 1 and 2.

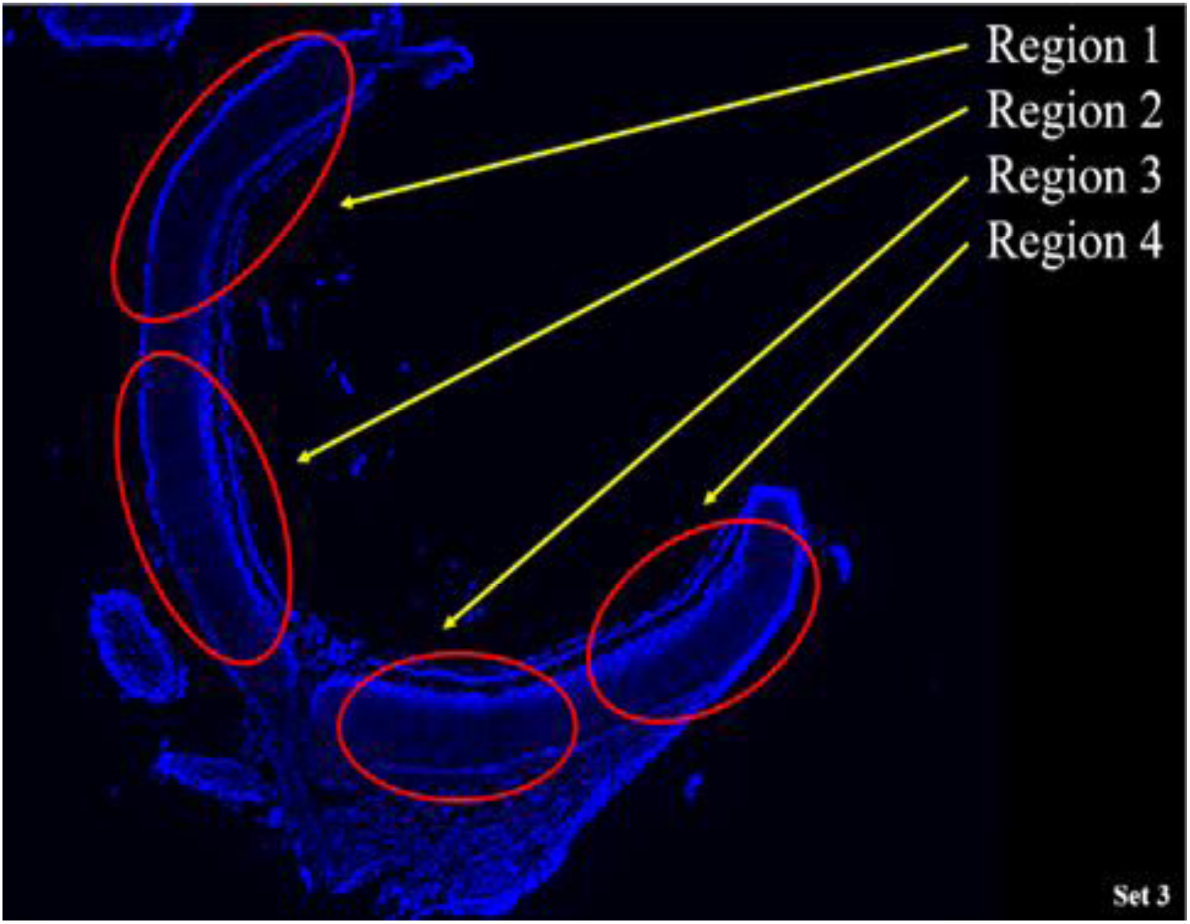

*Arr3* Microscope Imaging: The above image is a visual representation of each region in Set 3.

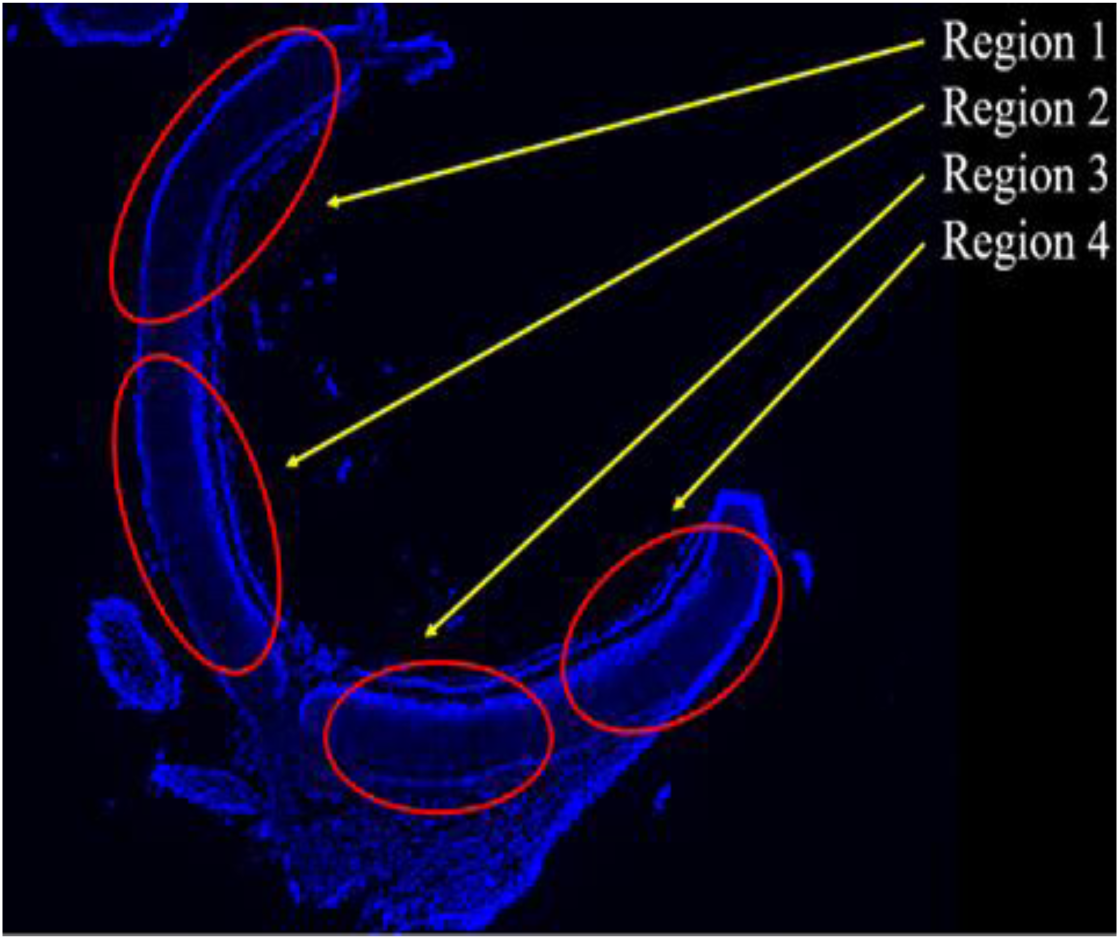

*Onecut1* Microscope Imaging: The above image is a visual representation of each region in all three sets.

#### Data Quantification

For PAX6 and BRN3A, no standardized rectangle was used over the imaged regions because the purpose was to count the total number of DAPI-stained cells in the GCL, and then calculate the percentage of positive PAX6 and BRN3A cells.

For *Onecut1*, *Glul*, and *p16*, ImageJ, a Java-based image processing system, was used to standardize the counting of the *Onecut1* gene. All images were rotated as close as possible to 0°, and a standardized rectangle was made over a specified region in the images of all three sets. *Onecut1* cells were counted in the INL and the GCL.

For *Arr3*, *RBPMS*, and *Nefl*, ImageJ was also used to standardize the counting of the *Arr3* gene. There were two standards in place for data quantification. For Standard 1, all images were rotated as close as possible to 0°. Then, a standardized rectangle was made over a specified region in the image. However, this standardized rectangle differed between sets because the Set 1 images were zoomed out 3x more than Sets 2 and 3. For Standard 2, all images were rotated as close as possible to 0°. No standardized rectangle was used because the purpose was to count the total number of *Arr3* cells and then calculate the percentage of *Arr3* cells on the border between the OPL and the ONL.

For all proteins and gene sets, including a larger area increased the sample size and ensured more accurate results.

## Statistical Analysis

Statistical methods are indicated in each figure.

---

**Supplementary Figure 1.**
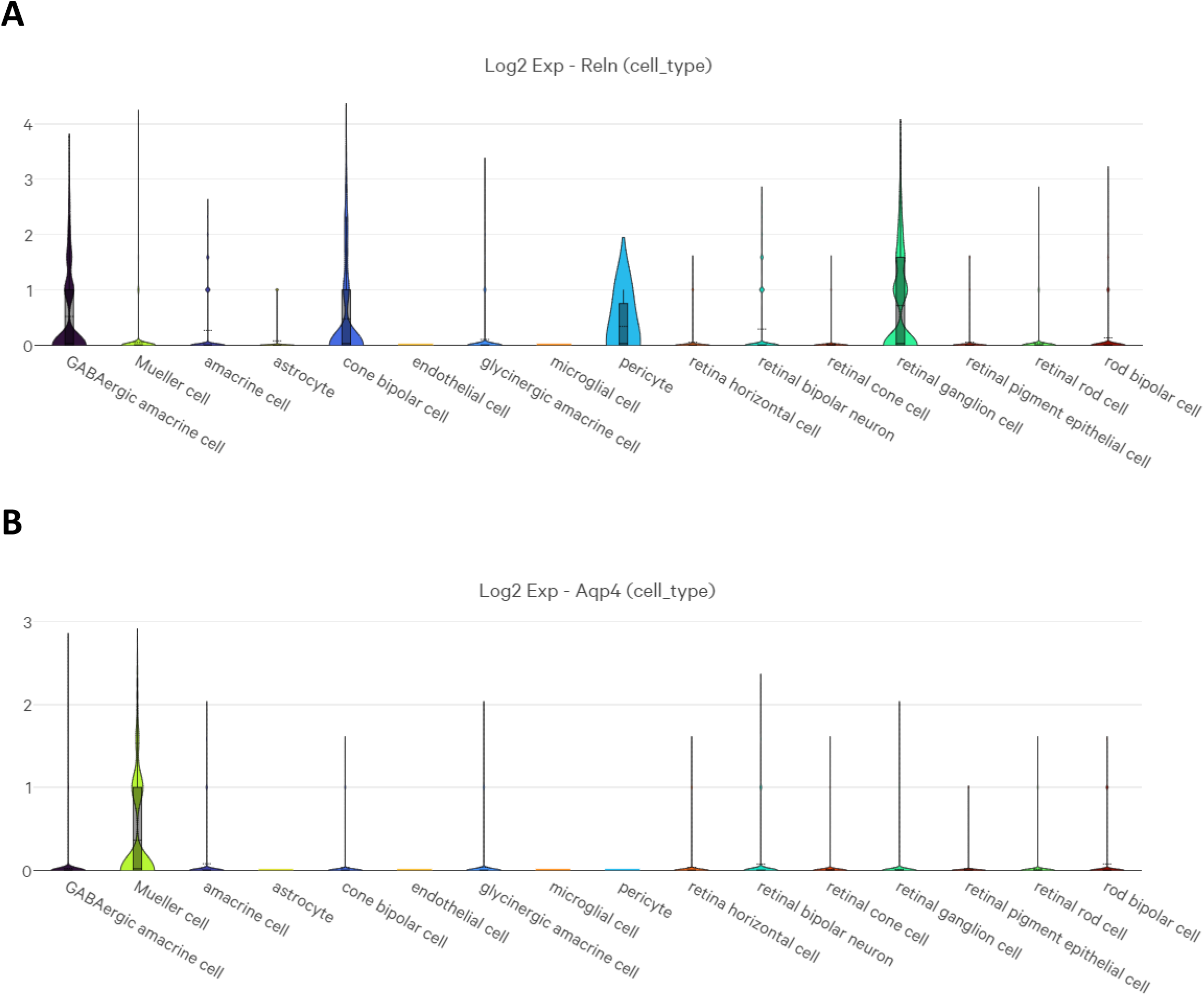
Cell-Type-Specific *Reln* Expression in the Adult Mouse Retina. **(A)** Violin plot showing cell-type specific expression of *Reln* in the snRNAseq dataset. **(B)** Violin plot showing expression of specific marker of Müller glia cells, *Aqp4* across annotated retinal cell types.

**Supplementary Figure 2.**
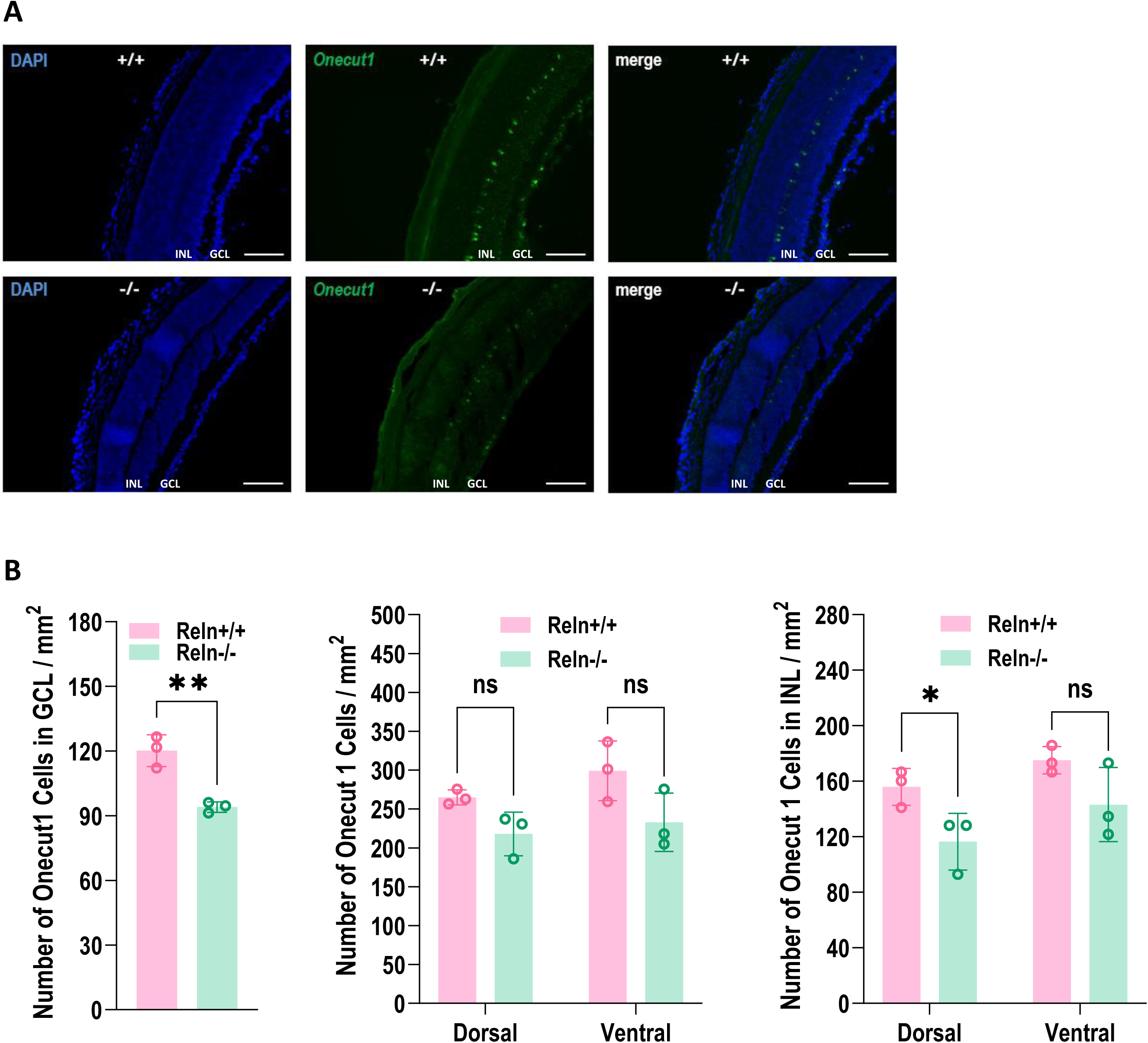
Regional Analysis of *Onecut1*⁺ Horizontal Cells in *Reln*⁻^/^⁻ Retina. **(A)** Representative dorsal retinal sections from *Reln*⁺^/^⁺ and *Reln*⁻^/^⁻ mice showing *Onecut1* with DAPI. Layers indicated: ONL, INL, GCL. Objective, 20×. Scale bars = 100 µm. **(B)** Quantification. Left: *Onecut1^+^* cells in the GCL (cells/mm²). Middle: Total *Onecut1*^+^ cells (cells/mm²) plotted separately for dorsal and ventral retina. Right: INL-restricted *Onecut1*⁺ cells (cells/mm²) by region. Bars show mean ± SEM; dots denote individual fields, averaged per animal for statistics (n = 6 fields from 3 animals per genotype). Significance: *p < 0.05, **p < 0.01 (unpaired two-tailed t-test). GCL = ganglion cell layer.

**Supplementary Figure 3.**
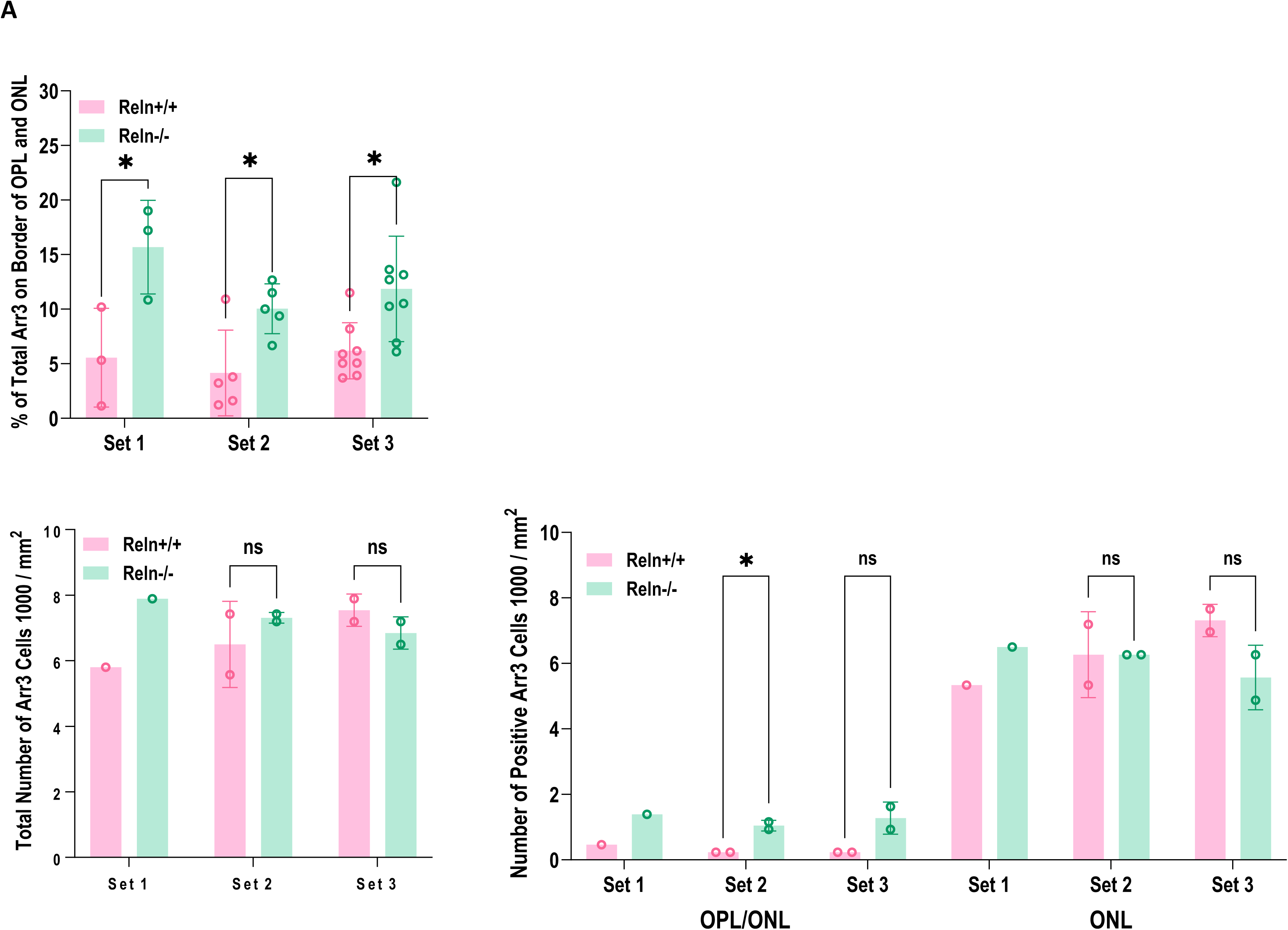
Reelin Deficiency Results in Stalled Cone Migration. **(A)** Top graph: Quantification of the percentage of *Arr3*+ cones across three independent biological replicate sets of *Reln*^+/+^ and *Reln*^−/−^ retinas. Bottom left: Total density of *Arr3*+ cones per mm² in both genotypes. Bottom right: Regional analysis shows number of positive *Arr3* per mm^2^ in thousands of cells in each set at both the border of the outer plexiform layer and outer nuclear layer and the outer nuclear layer only. Bars represent mean ± SEM; Significance: \**p* < 0.05, unpaired *t*-test.

## Notes

### Competing Interest Statement

The authors have declared no competing interest.

